# Region-Based Epigenetic Clock Design Improves RRBS-Based Age Prediction

**DOI:** 10.1101/2023.01.13.524017

**Authors:** Daniel J. Simpson, Qian Zhao, Nelly N. Olova, Jan Dabrowski, Xiaoxiao Xie, Eric Latorre Crespo, Tamir Chandra

## Abstract

Recent studies suggest that epigenetic rejuvenation can be achieved using drugs that mimic calorie restriction and techniques such as reprogramming induced rejuvenation. To effectively test rejuvenation *in vivo*, mouse models are the safest alternative. However, we have found that the recent epigenetic clocks developed for mouse reduced-representation bisulphite sequencing (RRBS) data have significantly poor performance when applied to external datasets. We show that the sites captured and the coverage of key CpGs required for age prediction vary greatly between datasets, which likely contributes to the lack of transferability in RRBS clocks. To mitigate these coverage issues in RRBS-based age prediction, we present two novel design strategies that use average methylation over large regions rather than individual CpGs, whereby regions are defined by sliding windows (e.g. 5 kb), or density-based clustering of CpGs. We observe improved correlation and error in our regional blood clocks (RegBCs) compared to published individual-CpG-based techniques when applied to external datasets. The RegBCs are also more robust when applied to low coverage data and detect a negative age acceleration in mice undergoing calorie restriction. Our RegBCs offer a proof of principle that age prediction of RRBS datasets can be improved by accounting for multiple CpGs over a region, which negates the lack of read depth currently hindering individual-CpG-based approaches.

## Introduction

DNA methylation (DNAm) based epigenetic clocks have become well established as predictors of chronological age (Bocklandt et al. 2011; Koch and Wagner 2011; Hannum et al. 2013; Horvath 2013; Weidner et al. 2014; Levine et al. 2018; Lu et al. 2019; Zhang et al. 2019). The predicted age generated by these clocks is referred to as epigenetic age (eAge) and the age acceleration (difference between eAge and chronological age, chAge) is associated with a number of disease states, conditions and all-cause mortality (Horvath et al. 2014; Horvath 2015; Horvath and Levine 2015; Horvath et al. 2016; Marioni et al. 2015; Chen et al. 2016; Simpkin et al. 2016; Maierhofer et al. 2017; Horvath et al. 2018; Lu et al. 2019; Martin-Herranz et al. 2019; Wu et al. 2019; Higgins-Chen et al. 2020; Fiorito et al. 2021; Jansen et al. 2021; Kim et al. 2022). As a result, eAge has become a widely used proxy to measure biological age, i.e. a measurement of age that may act as a better predictor of health and mortality than chronological age (Baker and Sprott 1988; Simpson and Chandra 2021).

Most human epigenetic clocks are based on methylation arrays, which offer precise and reproducible readouts of thousands of CpGs. Epigenetic age prediction in model organisms such as mouse enables the quantification of lifespan and healthspan interventions (Simpson and Chandra 2021; Simpson, Olova, and Chandra 2021). However, until recently, methylation arrays were restricted to human, and while arrays have become available for other mammals (Arneson et al. 2022; Zhou et al. 2022), they do not cover non-mammalian model organisms such as zebrafish. Reduced-representation bisulphite sequencing (RRBS) offers an alternative species-agnostic approach for the development of epigenetic clocks. For example, a number of mouse epigenetic age-predictors have been created using RRBS (Wang et al. 2017; Petkovich et al. 2017; Stubbs et al. 2017; Meer et al. 2018; Thompson et al. 2018; Trapp, Kerepesi, and Gladyshev 2021; see **S.Table 1** for a summary table of some of the main mouse RRBS clocks). In the original RRBS protocols, genomic DNA was enriched in CpG sites through enzymatic digestion with MspI (an enzyme that cuts DNA at CCGG sites, regardless of their methylation state; Meissner et al. 2005; Gu et al. 2011; Baheti et al. 2016) (**Fig. 1A**). This process is a cheaper alternative to whole genome bisulphite sequencing (WGBS) since it captures only ∼1% of the genome, achieving a higher relative depth of coverage (Meissner et al. 2008; Gu et al. 2011). However, RRBS datasets vary considerably in coverage and quality in comparison to the array format, which is highly standardised. This is possibly due to differences in library preparation protocols which could lead to inaccuracies when applying epigenetic clocks from one dataset to the next (Field et al. 2018; Thompson et al. 2018; Simpson and Chandra 2021).

**Figure 1:**
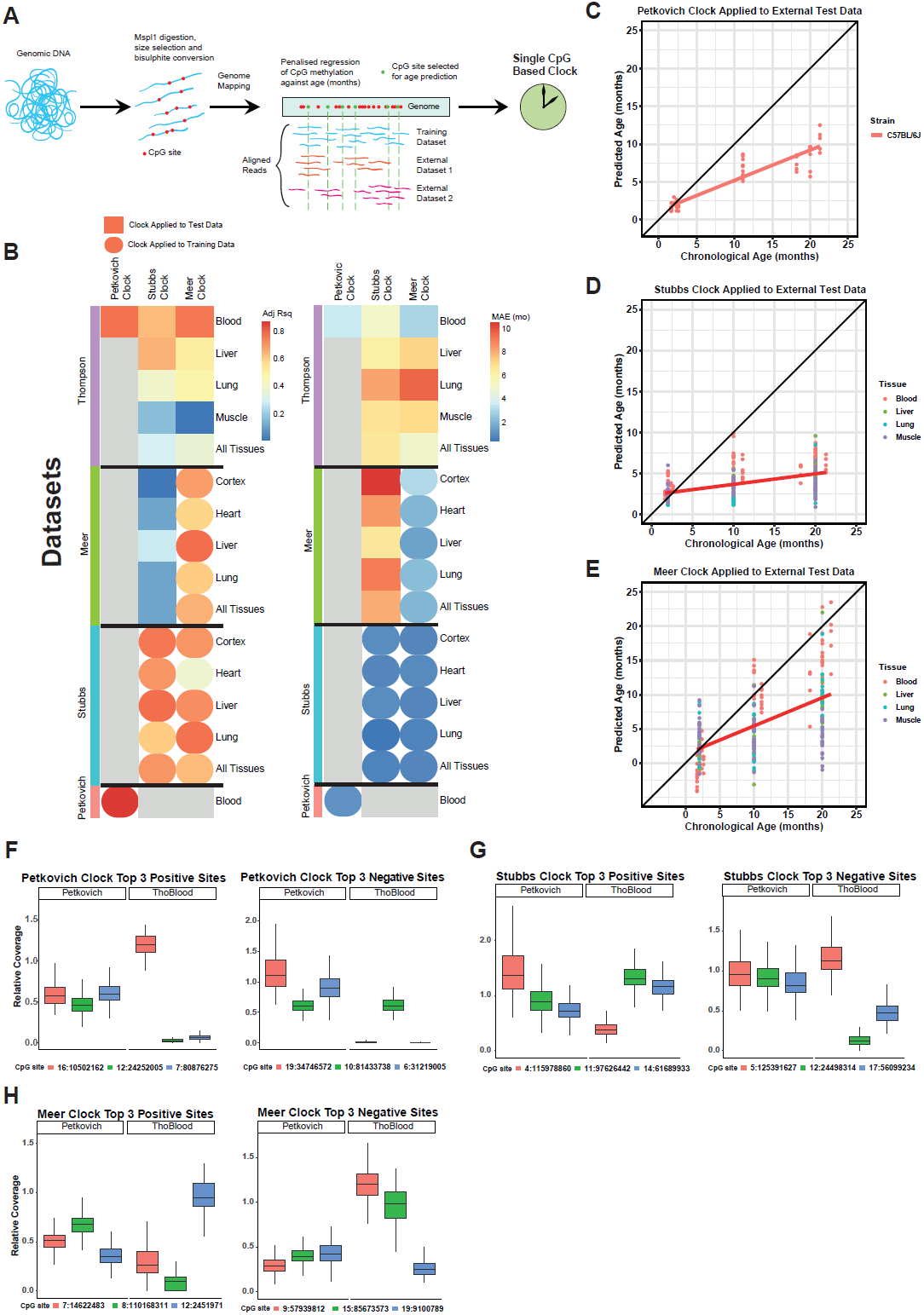
Mouse RRBS blood clocks are not as efficient when applied to external datasets (data independent of particular clock study). (**A**) Schematic of RRBS clock construction, with penalised regression conducted on methylation levels of single CpGs against age. (**Fig. 1B**) Adjusted R^2^ (left) and median absolute error (MAE, right) of various clocks applied to tissues from various datasets (Petkovich et al. 2017; Stubbs et al. 2017; Meer et al. 2018; Thompson et al. 2018). Round points represent clocks applied to training datasets, square points represent clocks applied to external datasets. (**C-E**) Petkovich (**C**), Stubbs (**D**) and Meer (**E**) mouse RRBS clocks applied to external Thompson multi-tissue dataset. (**F-G**) Relative coverage of the top three positive weighted sites (left) and top three negative weighted sites (right) of the Petkovich (**F**), Stubbs (**G**) and Meer (**H**) clocks in the Petkovich et al. 2017 and Thompson et al. 2018 datasets.

Here we show that the overlap of CpG sites captured in RRBS and their coverage differ between datasets, leading to poor transferability of clock algorithms between them. To overcome this limitation, we present a novel design strategy, generating clocks based not on individual CpG sites but on the average methylation level observed in contiguous regions. More precisely, we present two strategies, sliding windows and density-based clustering, to detect regions of interest that form the basis of our proposed regional clocks. These regional clocks outperform their individual-CpG-based counterparts both in accuracy and transferability. Furthermore, we show that they are more robust when applied to low-coverage, downsampled data and capture biological age.

## Results

### Uneven Coverage Contributes to RRBS Clocks Inaccuracy When Applied to External Datasets

We first analysed the performance of published mouse RRBS clocks against training data and data external to the study used to construct a particular clock. As expected, the clocks originating from Petkovich et al. 2017; Stubbs et al. 2017; Meer et al. 2018 (hereafter referred to as Petkovich, Stubbs and Meer clocks) had a high correlation and low error when applied to their training data (**Fig. 1B**). When these clocks were applied to external Thompson et al. 2018 C57BL/6 multiple tissue RRBS data, their overall performance was poor; median absolute error (MAE) was high (range 2.9-9.6 months) and R^2^ varied greatly (0.86-0.01) in age prediction (**Fig. 1C-E**). The best performance was achieved by the Petkovich and Meer clocks applied to blood (with R^2^ correlations of 0.86 and 0.85, respectively), however, their errors were still comparatively high (MAE = 3.5 months for Petkovitch, MAE = 2.9 months for Meer) versus their training data (MAE = 1.2 months for both clocks).

We hypothesized that the performance of RRBS clocks on external datasets is due, at least in part, to uneven coverage of CpGs captured across datasets from different studies (Field et al. 2018; Thompson et al. 2018; Simpson and Chandra 2021). To test whether uneven coverage contributes to the observed inaccuracies, we plotted from the Petkovich (used to train the Petkovich and Meer clocks) and Thompson datasets the relative coverage (normalised by total read count per sample) of the top three positive and negative weighted CpGs (as derived from their linear regression model) from the Petkovich, Stubbs and Meer clocks (**Fig. 1F-H**). We show that many of the top weighted CpGs required for age prediction in a given clock had a coverage close to zero in the Thompson external dataset, which may contribute to RRBS-based clocks not working as effectively on external datasets.

### Generating Regional Blood Epigenetic Clocks

It has been previously shown that proximal CpGs show correlated methylation behaviour (Lövkvist et al. 2016). We therefore explored whether taking the averaged CpG methylation value over a region (rather than from individual CpG positions) would overcome uneven coverage issues from RRBS data, resulting in more transferable age prediction (**Fig. 2A**). We used the mouse C57BL/6 whole blood training data from the Petkovich blood clock (aged 3-35 mo, training=129, cross-validation=12, sites with less than 5 reads filtered out; Petkovich et al. 2017) and segmented the mouse genome into various region sizes (1-9 kb) using sliding windows. For each window size, we calculated the averaged methylation levels from individual CpG sites within each window and conducted LASSO-penalised regression against chronological age to select optimal regions and weights for age prediction. Nine regional blood clocks (RegBCs) were generated (1-9 kb region sizes each, see **S.Table 3** for all RegBCs applied to training and cross-validation datasets). Considering R^2^ and MAE, we observed that the RegBCs’ performance on training and cross-validation data is relatively constant up to 6kb window size, after which the R^2^ in the cross-validation data decreases (**Fig. 2B**). Irrespective of the particular choice of window size, all RegBCs generated had R^2^ values higher than 0.93 and a MAE lower than 3 months when applied to training or cross-validation data, showing that regional clocks can perform at least as well as individual-CpG-value-based predictors. To investigate the extended power of regional clocks, we then proceeded to test the transferability of this new model to unseen data. For this we chose to test on an external blood dataset (Thompson et al. 2018) the 5kb window regional blood clock (5kb RegBC) as it has the smallest difference in error and R^2^ between the training and cross-validation dataset. The 5kb RegBC showed a higher correlation and lower error (R^2^=0.91, MAE=3.38, **Fig. 2C**) than the blood-specific Petkovich clock trained on individual CpGs (R^2^=0.86, MAE=3.52, **Fig. 1C**). The 5kb RegBC also has a higher R^2^ than the Meer clock applied on the Thompson data, albeit the Meer clock has a lower MAE (R^2^=0.85, MAE=2.94, **Fig. 1E**). The 5kb RegBC has 24 negatively weighted regions (regions that hypomethylate with age) and 11 positively weighted regions (regions that hypermethylate with age) (**S.Table 4**). A total of 1978 CpGs are present in the 35 total regions of the 5kb RegBC, with a mean of 57 CpGs per region.

**Figure 2:**
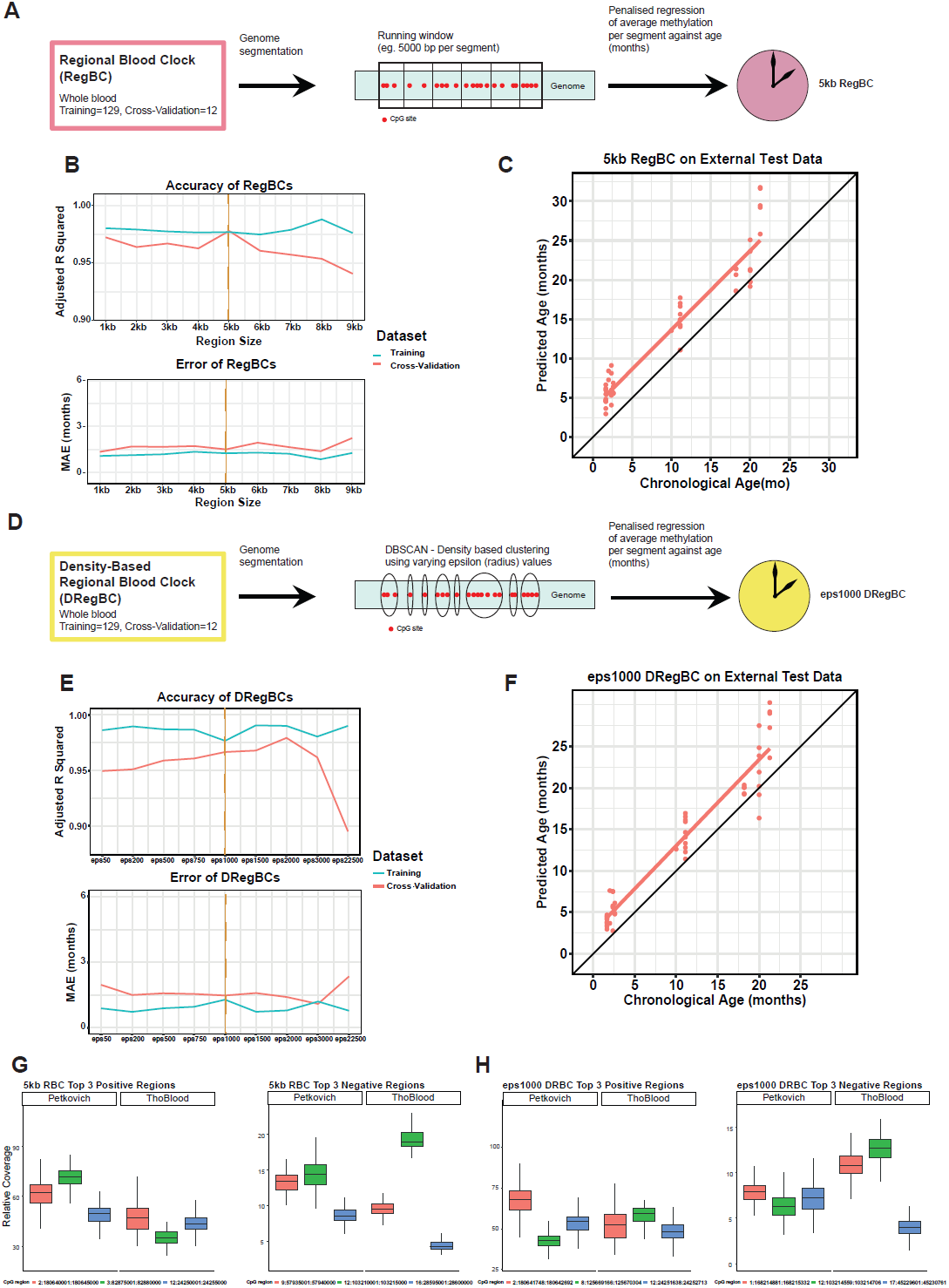
5 kb genome window size and density-based clustering provide transferable age prediction. (**A**) Graphical overview of regional blood clock (RegBC) creation. A running window generator was applied to segment the mouse genome into various sizes (eg. 5kb). Segments for each particular window size were trained against age (months, mo) using penalised regression, resulting in a given clock for a particular window size. (**B**) Nine RegBCs were generated (1-9 kb region sizes). Correlation (Adjusted R^2^ of epigenetic age correlated with chronological age, **top**) and error (MAE, **bottom**) of each RegBC (x-axis) applied to a particular dataset. Blue and red lines are internal training and cross-validation datasets respectively (Petkovich et al. 2017). Orange vertical dashed line highlights the 5kb RegBC as the best clock selected and taken forward. (**C**) 5kb RegBC applied to Thompson et al. 2018 external dataset. (**D**) Graphical overview of density-based regional blood clock (DRegBC) creation. The density-based spatial clustering of applications with noise (DBSCAN) algorithm was applied to the CpGs in the Petkovich et al. 2017 blood training data to group them based on distance. Various epsilon (eps) values were used to govern the distance (in base pairs) between each CpG to form a cluster. For each eps value, the resulting clusters were then trained against age using penalised regression, resulting in a given clock. (**E**) Nine eps values varying from 50 to 22500 were used to construct DRegBCs. Correlation (Adjusted R^2^ of epigenetic age correlated with chronological age, **top**) and error (MAE, **bottom**) of each DRegBC (x-axis) applied to a particular dataset (colours represent the same datasets as in **B**). Orange vertical dashed line highlights the eps1000 DRegBC as the best clock selected and taken forward. (**F**) eps1000 DRegBC applied to Thompson et al. 2018 external dataset. (**G**) Relative coverage of the top three positive weighted sites (left) and top three negative weighted sites (right) of the 5kb RegBC in the Petkovich et al. 2017 training data and Thompson et al. 2018 external dataset. (**H**) Relative coverage of the top three positive weighted sites (left) and top three negative weighted sites (right) of the eps1000 DRegBC in the Petkovich et al. 2017 training data and Thompson et al. 2018 external dataset.

Although the sliding window approach is widely used for whole genome segmentation, groups of CpGs with correlated methylation values could easily get separated into different regions, which may hinder age prediction. Therefore, we tested whether RegBCs built using CpGs clustered based on proximity to each other would improve age prediction. For this, we applied the density-based spatial clustering of applications with noise (DBSCAN) algorithm to the CpGs covered in the Petkovich RRBS blood training data (sites with fewer than 5 reads were filtered out) (Ester et al. 1996). To find clusters, we used the distance (base pairs) between CpG sites and required at least 5 CpG sites to form a cluster or region. DBSCAN then discovers regions of clustered CpG sites that are separated by a minimum distance or radius (epsilon value, eps), see **Fig. 2D**. We created regions of CpGs using varying eps values between 50 and 22500 bp. For each eps value, we again conducted LASSO-penalised regression against chronological age to select the optimal regions and weights to predict age. Nine density-based regional blood clocks (DRegBCs) were constructed with varying eps values (**S.Table 3**). All DRegBCs had R^2^ values exceeding 0.9 and an MAE less than 3 months when applied to the training and cross-validation datasets (**Fig. 2E**). The eps1000 DRegBC had the smallest difference between training and cross-validation datasets in terms of both R^2^ and MAE (with the exception of the eps2000 DRegBC which had a lower MAE on its cross-validation test). We applied it to the external Thompson blood dataset, where it outperformed the Petkovich and Meer clocks based on individual CpGs (R^2^=0.92, MAE=2.71, **Fig. 2F**). The mean region size found in the eps1000 DRegBC is 623 bp, with a mean of 55 CpGs present per region. There are 33 regions in total, with 21 negatively weighted and 12 positively weighted.

To test whether our approach reduced coverage differences, we plotted the coverage for top weighted CpGs between data sets. We found that the relative coverage of the top three hyper/hypomethylating regions in the 5kb RegBC and eps1000 DRegBC in the Petkovich and Thompson datasets is largely maintained using the regional approach (**Fig. 2 I,J**). Hence, taking the averaged value of CpG methylation over regions appears to improve relative proportions of reads for clock components between datasets compared to individual CpG approaches (**Fig. 1 F-H**), which may contribute to the improved age prediction in external data sets.

Between the 5kb RegBC and eps1000 DRegBC, 21 5kb RegBC regions overlap with 22 eps1000 DRegBC regions (with one DRegBC site overlapping with two RegBC windows) (**S.Table 4**). These overlapping regions (many of which were top weighted in both clocks) were also associated with various genes (*Tcfl5, Map10, Smarca5-ps, Rbm46, Aldh1a2* and *Gm21297* in hypermethylating regions and *Prima1, Fgf12, Eml4, Qrfp, Hpn, Prr13* and *Ntn1* in hypomethylating regions). We also found a number of CpGs from the Petkovich and Meer clocks present in our 5kb RegBC and eps1000 DRegBC regions, as expected from these clocks being created using the same training data. 16 Petkovich clock CpGs overlap with 6 regions of 5kb RegBC, while 15 overlap with 5 regions of the eps1000 DRegBC, all of which are regions shared by both clocks except one site/region unique to the 5kb clock. The Meer clock is more represented in both clocks, with 23 CpGs in 7 of the 5kb RegBC regions, while 19 Meer clock CpGs are overlapping with 5 eps1000 DRegBC regions, all of which are shared with the 5kb RegBC. 11 Meer CpG sites were present in one particular region in the eps1000 DRegBC, 2:164167686:164169038, which is also in a 5kb RegBC region. This suggests that, as expected, the eps1000 DRegBC finds more targeted regions than the 5kb RegBC clock.

### Biological Topography and Relevance of Regional Blood Clocks

There is a large degree of overlap between the 5kb RegBC and the eps1000 DRegBC, however, regions captured by the eps1000 DRegBC are on average smaller. This would suggest the DBSCAN method identifies more discrete regions that might allow us to capture some degree of underlying biological relevance. We first plotted the genomic GC content percentage for each region, ordered by the slope of methylation change with age. Globally, AT-rich regions tend to be heterochromatic and constitutive lamina-associated domains (Saccone, Federico, and Bernardi 2002; Federico et al. 2004; Gilbert et al. 2004; Bolzer et al. 2005; Federico et al. 2006; Meuleman et al. 2013; Chandra et al. 2015). We observed a strong correlation (R = 0.70) between average GC content and hypo/hypermethylation gradient (**Fig. 3A,B**). Hypermethylating regions show an average GC content as high as 80%, while hypomethylating regions were as low as 40% (**Fig. 3A**). This suggests that age-dependent hypomethylation is associated with heterochromatic areas, consistent with previous observations (Bollati et al. 2009). In contrast, CpG islands (CGIs) were almost exclusively found in hypermethylating regions (**Fig. 3B**), consistent with previous studies that show CGIs tend to gain methylation with age (Christensen et al. 2009; Heyn et al. 2012; Johansson, Enroth, and Gyllensten 2013; Gopalan et al. 2017; Slieker et al. 2018).

**Figure 3:**
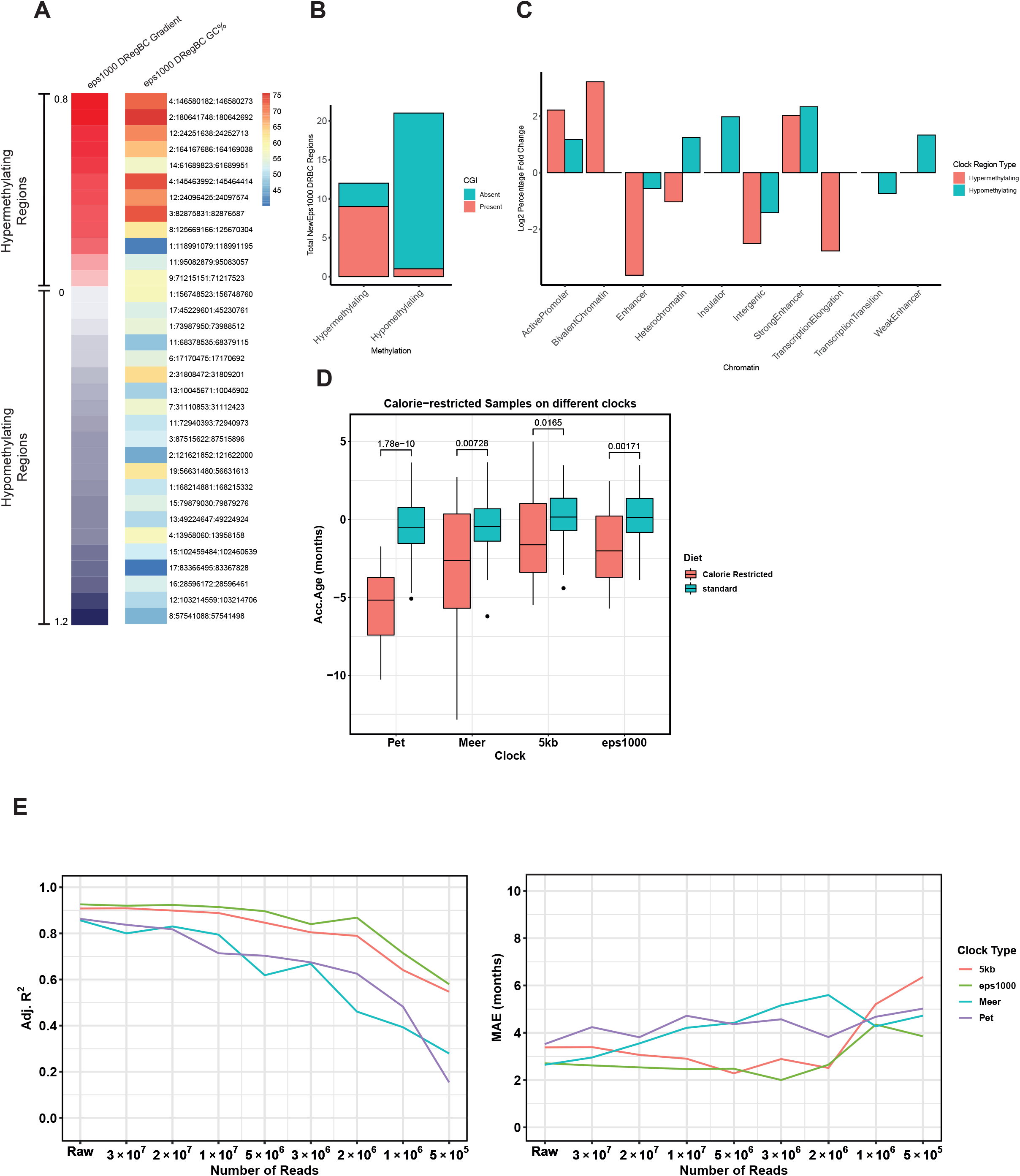
Chromatin structure and biological age prediction of eps1000 DRegBC. (**A**) Percentage GC content of eps1000 DRegBC regions ordered by the gradient of methylation change with age. (**B**) Number of eps1000 DRegBC regions overlapping with CpG islands (CGIs) in hypermethylating and hypomethylating regions. (**C**) Log2 fold change of ChromHMM states proportions in eps1000 DRegBC hypermethylating or hypomethylating regions compared to genomic proportions. (**D**) Petkovich, Meer, 5kb RegBC and eps1000 DRegBC applied to calorie restricted and control samples. (**E**) Correlation (adjusted R^2^ of epigenetic age correlated with chronological age, left) and error (MAE, right) of 5kb RegBC, eps1000 DRegBC, Petkovich and Meer clocks applied to downsampled Thompson et al. 2018 external dataset.

To annotate the eps1000 regions in more detail, we looked at their overlap with ChromHMM chromatin states (**Fig. 3C**; see **S.Table 5**) (Ernst and Kellis 2012; Pintacuda et al. 2017). We calculated the percentage bp of each ChromHMM state found within each eps1000 DRegBC window and calculated the log2 fold difference between chromatin proportions in the clock regions and the genome (see “Annotating eps1000 DRegBC with ChromHMM Labels and Plotting Log Fold Change” in Methods). Compared to the genome, the eps1000 DRegBC regions had a lower proportion of intergenic regions and a higher proportion of active promoters. The eps1000 DRegBC had a higher proportion of heterochromatin in hypomethylating regions compared to hypermethylating regions, which is consistent with the GC proportions in **Fig. 3B** and previous findings (Bollati et al. 2009). Bivalent chromatin occurs at a higher proportion in hypermethylating than hypomethylating regions, which is consistent with hypermethylation of bivalent chromatin as a hallmark of ageing and with previous clock studies (Rakyan et al. 2010; Horvath 2013).

### Regional Blood Clocks Capture Biological Age

To investigate whether the 5kb RegBC and eps1000 DRegBC can capture biological age, we applied each clock, as well as the Petkovich and Meer clocks, to calorie restricted (CR) samples (**Fig. 3D**). These CR samples were not included in the training and show a significant age deceleration for both 5kb RegBC and eps1000 DRegBC (median error = -1.6 and -2.0 months, respectively). The Petkovich clock showed a greater age deceleration of -5.1 mo, while the Meer clock has an age deceleration of -2.6 mo, similar to the eps1000 DRegBC.

### Regional Blood Clocks Are More Robust When Applied to LowCoverage Data

We next explored whether low coverage data could benefit from regional clock designs resulting in more robust age-prediction upon downsampling. We step-wise randomly downsampled the Thompson blood external dataset from raw (40 million reads) to 500,000 reads, and applied the 5kb RegBC, eps1000 DRegBC, Petkovich and Meer clocks to each level of downsampling (**Fig. 3E**). Both regional-based clocks maintain an R^2^ above 0.75 up to 2 million reads, while at this point, the Petkovich clock declined in correlation to ∼0.46. At 500,000 reads, both regional-based clocks maintain a correlation above 0.55, while the Petkovich and Meer clocks are below 0.4. Both regional-based clocks have an MAE lower than 4 months until 1 million reads. Interestingly, random downsampling between 30 million and 3 million reads appears to lower error as both regional clocks have an error below 3 mo, with the eps1000 DRegBC as low as 2 months at 3 million reads. At 500,000 reads, the eps1000 DRegBC error increases to ∼4 mo, while the 5kb RegBC increases to ∼6.5 mo. The Meer and Petkovich clocks by comparison are more stable with an error of ∼5 mo, however, the eps1000 DRegBC remains the most accurate with the lowest error and highest correlation at 500,000 reads.

## Discussion

Age prediction utilising RRBS data has proven difficult due to the uneven coverage of CpG sites captured between different experiments. We have shown that a regional approach, based either on set window sizes or dynamically allocating windows based on CpG density, can improve the transferability of the age predictor when applied to external datasets as well as robustness when applied to low coverage data. Using samples from the same dataset used to create the Petkovich blood clock, our RegBCs outperform individual-CpG-value-based clocks (Petkovich and Meer clocks) when tested on unseen external blood datasets. However, the main limitation of this study is the limited number of suitable RRBS datasets available for training and testing our approach.

An important characteristic of epigenetic age predictors is their ability to capture biological perturbations of ageing. The regional blood clocks were able to detect a negative age acceleration in calorie-restricted mice. Our method was also able to offer more biological context by revealing the chromatin topography of regions useful for age prediction. Many regions were found to be shared by both the 5kb RegBC and eps1000 DRegBC, with CpGs from published clocks also present. In these regions shared by both clocks, it is likely that the age-related methylation changes fall in the eps1000 DRegBC regions specifically, since all the window sizes for the eps1000 DRegBC are smaller than the 5kb windows. Indeed, one eps1000 DRegBC region was present in two 5 kb RegBC regions, hence a cluster of CpGs with correlated methylation values was split up due to the sliding window approach, which may inhibit age prediction and biological interpretation. In addition, the eps1000 DRegBC was more accurate in predicting age than the 5kb RegBC. Therefore, a cluster-based approach for assembling regions rather than a windowed approach might be more effective for both predicting age and gleaning biologically significant results. For example, a high proportion of bivalent (poised) chromatin was found in regions that hypermethylate with age in the eps1000 DRegBC, which is a known hallmark of ageing (Rakyan et al. 2010). We also found enrichment of heterochromatin in hypomethylating regions, which is consistent with previous studies (Bollati et al. 2009). In addition, the majority of hypermethylating regions in our regional clocks were overlapping with CpG islands. This is, however, expected as CpG islands are commonly hypomethylated and, therefore, can only gain in methylation with age, which is a well documented phenomenon (Heyn et al. 2012; Johansson, Enroth, and Gyllensten 2013; Gopalan et al. 2017; Slieker et al. 2018). Indeed, CpG islands bound by PRC2 *de novo* methylate with age in certain tissue types (Maegawa et al. 2010; Klutstein et al. 2017) and methylation gain in these regions can be used to track ageing (Moqri et al. 2022). It is therefore possible that the regions found in our clock may be similar regions where methylation is dysregulated with age, however, mechanistic studies are required to confirm any causal role in ageing (Moqri et al. 2022).

Targeted methods have been developed for mice to reduce the cost that comes with RRBS and WGBS. Recently, the Wagner lab developed a 3 CpG and a 15 CpG clock for multiple platforms (pyrosequencing, droplet digital PCR and barcoded bisulphite amplicon sequencing), which offered accurate age prediction at few CpGs for relatively low cost (Han et al. 2018; Han et al. 2020). Techniques such as this offer certain benefits for age prediction since they have a higher coverage at a small selection of CpGs, whereas RRBS experiments vary in coverage necessary for the CpG-specific approach for age prediction (**Fig. 1E**,**F**). Our regional clocks offer a proof of principle that RRBS-based age prediction can be made more effective by accounting for multiple CpGs over a region and negating the lack of read depth at key CpGs that would otherwise hinder single CpG-based clocks. We show that regional clocks outperformed individual-CpG-based clocks when applied to downsampled datasets, showing that the averaged methylation over regions is able to compensate for low coverage, offering a way to generate clocks based on shallower sequencing. This principle could be applied to RRBS datasets (and possibly WGBS) from other model organisms to develop age predictors. In addition, our technique has highlighted key regions that could be specifically targeted for accurate age prediction in mouse blood.

## Methods

### Data Collection

The following mouse RRBS datasets were downloaded from the NCBI Gene Expression Omnibus (GEO); GSE80672 (141 C57 mouse and 20 calorie restricted C57 mouse; Petkovich et al. 2017), GSE120137 (n=548; Thompson et al. 2018), GSE93957 (n=62; Stubbs et al. 2017), and GSE121141 (n=81; Meer et al. 2018). The annotation information (metadata) of the sequencing samples, such as age and tissue, was downloaded from the NCBI SRA Run Selector. The unit of the chronological age is different among datasets, including month and week. For the convenience of subsequent calculations, all ages in weeks were converted to age in months. The conversion formula is:

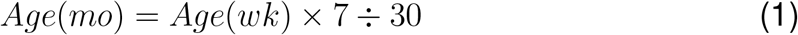

where *Age*(*mo*) is an age in months, *Age*(*wk*) is an age in weeks, 7 means 7 days per week, and 30 means average 30 days per month.

### Data Processing

The following processes were implemented via bash scripts on Eddie, a high performance computing cluster with a Linux-based operating system provided by the University of Edinburgh.

Read adaptors were removed by TrimGalore (ver 0.5.0, –rrbs) which combines FastQC (ver 0.11.4) and cutadapt (ver 1.9.1; Krueger 2012). The following processing steps were conducted with Bismark (ver 0.18.1), a utility designed to process and map BS-seq data using packages such as bowtie2 (mapping, ver 2.2.6) and samtools (read processing, ver 1.6; Krueger and Andrews 2011):

1. A bisulphite converted version of GRCm38.p6 reference genome was created by the Bismark genome preparation module (Krueger 2016).
2. The trimmed reads were aligned to the bisulphite converted genome with Bismark (settings: –multicore 2 –phred33-quals -N 0 -L 20).
3. Certain samples in the Petkovich dataset were processed twice on different flow cells. Duplicate sample BAM files were merged post mapping with Rsamtools (ver 1.36.1; Morgan M et al. 2022).
4. The Bismark methylation extractor module then obtained from the BAM files (generated in the second step) the mapped reads and methylation count information (CpG, CHG and CHH contexts). CHG and CHH sites were removed.
5. The bismark2bedGraph module was used to extract the methylation information from CpG context files and convert them to cov files (bismark.cov.gz) which contain six columns: (1) chromosome number; (2) start position; (3) end position; (4) percentage of methylated reads in total reads (β score); (5) the number of methylated reads; (6) the number of unmethylated reads (Krueger 2016).

### Coverage Assessment

To compare whether there was a difference in coverage between all downloaded datasets (see “Methods: Data Collection” for list of datasets), the number of reads from the cov files was extracted using the “Read Count Quantitation” pipeline in SeqMonk (ver 1.48.0; Andrews 2007):

1. Create a new project. “File” *→* “New project” *→* select genome GRCm38 v100 *→* click “Start New Project”.
2. Load RRBS data. “File” *→* “Import Data” *→* select “Bismark(Cov)” *→* select cov files in the new window.
3. Add annotation (for eps clock). “File” *→* “Import Annotation” *→* select “Text(Generic)”
4. Define a probe.
  - For kb clock, “Data” *→* “Define Probes” *→* select “Running Window Generator” *→* set probe size.
  - For eps clock, “Data” *→* “Define Probes” *→* select “Feature Probe Generator” *→* select the annotation imported by the previous step (do not select “remove exact duplicate”).
  - For single-site clocks, “Data” *→* “Define Probes” *→* select “Read Position Probe Generator” *→* select samples.
5. Define quantitation.
  - For coverage, “Data” *→* “Quantitate Existing Probes” *→* select “Read Count Quantitation” (do not choose “Correct for total read count” and “Log-Transform Count”).
  - For methylation value, “Data” *→* “Quantitation Pipelines” *→* “Bisulphite methylation over features” *→* “Run Pipeline”
6. Generate reports which contain coverage or methylation value. “Reports” *→* “Annotated Probe Report”

The coverage of Petkovich, Stubbs, Meer and Thompson blood data was extracted and organised into a matrix, which was loaded into R (ver 4.1.1). The relative coverage was calculated as follows:

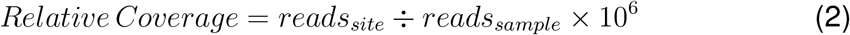

Representative sites (top 3 negatively and positively related to age) were selected based on their weight in a given clock. Each column represented a CpG site (or region of CpGs in the case of the regional clocks). Coverage of clock sites/regions in different datasets was plotted using ggplot2 (ver. 3.3.3; Wickham 2016) in R (**Fig. 1F-H, Fig. 2G,H**).

### Applying Published BS-seq Clocks to Various Datasets

The Petkovich blood clock and two multi-tissue (Meer and Stubbs) mouse BS-seq clocks were tested on various datasets. For each clock, the required CpG sites were extracted from a given dataset that the particular clock would be applied to (**S.Table 1**). Each clock was ran according to scripts and instructions from their respective publications, e.g. the toRun Imputation. An R script provided by Stubbs et al. (2017) was used to run their clock (Wang et al. 2017; Petkovich et al. 2017; Stubbs et al. 2017; Meer et al. 2018; Thompson et al. 2018).

To evaluate the results, the adjusted R squared (adj. R^2^) and median absolute error (MAE) were calculated, and linear regression lines were illustrated with ggplot2 in R (Wickham 2016). The formula for adj. R^2^ is as follows:

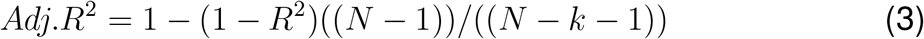

where *N* represents the number of samples, and *k* represents the number of predictors (Miles 2014). MAE is the median absolute error between epigenetic age and the chronological age, where if a test dataset has an MAE of X, then the age acceleration will differ by less than X in 50% of the samples (Horvath 2013). The formula is as follows:

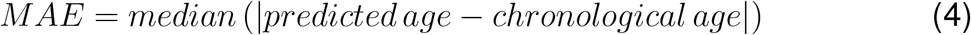

### Genome Segmentation

Before constructing a regional epigenetic clock, we filtered out any CpGs with fewer than 5 reads. Genome segmentation with set window sizes (e.g. 5kb) was conducted using SeqMonk (ver 1.47.0; Andrews 2007), an interactive desktop application with a suite of tools for analysing mapped genomic data. GRCm38 v100 was selected as the reference genome. Next, the filtered cov files were imported into SeqMonk, “Define Probe” was opened from the drop-down window of “Data”, and then “Running Windows Generator” was selected to set the window size (probe size) and step size. Window size and step size were set as equivalent, e.g. 5kb for both to avoid any overlapping windows. After setting the probe, “Bisulphite methylation over features” was selected from “Quantitation Pipelines” to calculate the average methylation level per window/region. The formula of the average methylation level per region is:

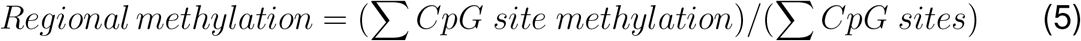

where *CpG site methylation* is the mean methylation of an individual CpG site within a given region. This method was preferred than to take mean methylated reads and dividing by total unmethylated reads in a given region as this does not account for coverage bias due to a potential small number of highly methylated positions (Olova et al. 2018). There could also be very few reads, which would also result in an unreliable mean methylation call (Andrews 2018). The results were exported as a counts matrix text file, where samples are columns and rows are genomic regions. Each dataset (see “Methods: Data Collection”) was segmented for all window sizes (1-9kb) and were used for either training or testing the regional clocks.

For construction of the density-based RegBCs (DRegBCs), the locations of single CpG sites in the Petkovich training data was extracted using the “Read Position Probe Generator” function in SeqMonk to create probes for single CpGs. The coordinates were loaded into R and a custom for loop applied the dbscan function (from the dbscan package (Hahsler, Piekenbrock, and Doran 2019)) for various epsilon (eps) values (50, 200, 500, 1000, 3000, 22500) to each chromosome. In the case of constructing a regional clock, the eps value is the maximum number of base pairs (distance) between each point to form a cluster. Minimum points to call a cluster was left at default (5 points, i.e. 5 CpGs). The mean methylation for each cluster was calculated in SeqMonk in the same manner as the RegBCs above.

### Regional Epigenetic Clock Construction Using LASSO

Each text file of region counts per region clock iteration (generated by SeqMonk) was loaded into R. All percentage methylation values for each region were divided by 100, so that 0-1 would represent 0% to 100% respectively. Regions with NaN (ie. no reads recorded) were removed). Least Absolute Shrinkage and Selection Operator (LASSO) regression model from the glmnet package in R (ver 4.1.1) (Friedman, Hastie, and Tibshirani 2010) was applied. LASSO is a penalised regression model which selects the best CpG regions that correlate with age in a linear regression model:

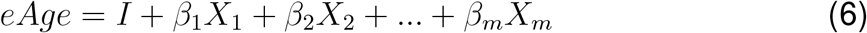

where *eAge* represents the epigenetic age,*I* is the intercept value where the linear model would meet the y-axis, *β* represents the average methylation score of a particular genomic region and *X* is the coefficient/weight (Hepp et al. 2016). We ran a 100-fold cross validation (using the cv.glmnet function) for each clock, saved the lowest lambda value determined and used this value when running the LASSO model for each clock (using the glmnet function; other than lambda, all other terms were default). A list of regions, their coefficients, and an intercept value were then generated.

Nine regional blood clocks (RegBCs, (1-9 kb region sizes) and nine densitybased RegBCs (DRegBCs, varying eps values) were generated **S.Table 3**). Whole blood from GSE80672 (training=129, cross-validation test=12; Petkovich et al. 2017) was used to construct the RegBCs. The Petkovich blood training dataset was randomly partitioned into 90% training and 10 % cross-validation using the caret package (ver 6.0-86; Kuhn 2020) in R.

Each clock was applied to a particular dataset using a for loop written in R. It extracts each required CpG region (generated from LASSO penalised regression) from each sample in the counts matrix (generated from SeqMonk) and multiplies it with its corresponding weight (generated from the LASSO penalised regression) as according to equation (6). Each iteration of the RegBC was applied to the blood training and cross-validation datasets (adj. R^2^ values were plotted in **Fig. 2B**, left panel) and the Thompson et al. (2018) blood samples (GSE120137) (adj. R^2^ and MAE values were recorded in **S.Table 3**). Any values that were missing in the cross-validation or external datasets were imputed as zero.

### Regional Epigenetic Clock Applied To Calorie Restricted Samples

RRBS data from calorie restricted C57 mice (CR, n=20) was downloaded. For RegBC age prediction, genome was segmented and average methylation was calculcated per region as per previous steps for either 5kb RegBC or eps1000 DRegBC. 5kb RegBC, eps1000 DRegBC, Petkovich blood clock, and Meer clock were applied to CR data aged 10-27 months as per previous steps (see Methods: “Applying Published BS-seq Clocks to Various Datasets”). The predicted age of CR data were compared to a control group (n=74) which is C57 mice aged from 10 to 28 months sampled from the Petkovich blood data (training + crossvalidation test = 141). Age acceleration was then calculated as predicted age minus chronological age. ggboxplot (ggpubr ver 0.5.0) was used to plot age acceleration. Wilcoxon test were used to compare the differences between the two groups. The median of the age acceleration was calculated in R and reported as the median error.

### Annotation of Regional Blood Clocks with Published Blood Clock Sites, GC Counts and Other Metadata

Annotation files for the 5kb RegBC and eps1000 DRegBC were initially created using SeqMonk and collated in **S.Table 4**. For example, to annotate the 5kb RegBC with another clock, such as the Petkovich clock, the “Feature Probe Generator” in SeqMonk was used to generate probes based on each of the Petokvitch clock sites. “Bisulphite methylation over features” was then calculated (these values were not used, but quantitation was required to create an annotated probe report). An annotated probe report was created where each Petkovich site was annotated with an overlapping 5kb RegBC window. In R, the results were collated to count the number of Petkovich CpGs that are present in each 5kb RegBC region. These counts were then added to **S.Table 4**. This process was then repeated for the Meer clock. Similarly, overlapping eps1000 DRegBC regions and CpG islands (mm10 coordinates from Olova et al. 2018, originally from Illingworth et al. 2010) were mapped and processed in the same manner but instead of counts, names of overlapping regions were pasted directly into **S.Table 4** for each 5kb RegBC region. This entire process was also done in the same way to annotate the eps1000 DRegBC. Total CGIs present in hyper- and hypomethylating regions were plotted using ggplot2.

Total base content for each RegBC probe was calculated with a custom perl script from https://github.com/NellyOlova/BS_bias (Olova et al. 2018). GC percentage per clock region was then calculated as total G bases plus C bases divided by region length. GC percentage for each eps1000 DRegBC region was then plotted using pheatmap (ver 1.0.12) in R, ordered by methylation change (gradient) with age (calculated using the lm function in R). Methylation gradient was also plotted with pheatmap. Hyper- and hypomethylating regions were defined as any region with a positive or negative clock coefficient respectively.

### Annotating eps1000 DRegBC with ChromHMM Labels and Plotting Log Fold Change

mm10 ChromHMM labels were downloaded from https://github.com/guifengwei/ChromHMM_mESC_mm10 (dense annotated bed file used) (Pintacuda et al. 2017). The bed file was loaded as an annotation track in Seqmonk. For our training data, probes were created based on the mESC bed file annotations using the “Feature Probe Generator” function. “Bisulphite methylation over features” was then calculated (these values were not used, but quantitation was required to create an annotated probe report). An annotated probe report was created, where the mESC ChromHMM probes were labelled with either overlapping 5kb RegBC regions and eps1000 DRegBC regions. To calculate the number of base pairs each chromatin state was present within each eps1000 DRegBC region, the start site in bp for a given ChromHMM annotation was subtracted from the start site of the corresponding eps1000 DRegBC window. This gave the “starting flank”, i.e. the number of bp within the ChromHMM annotation a particular clock region starts. The “ending flank” of the same ChromHMM annotation/eps1000 DRegBC region was then calculated as the eps1000 region end coordinate subtracted from the ChromHMM end coordinate. Any starting or ending flanks that had values lower than 0 (i.e. negative values) were therefore the number of bp a ChromHMM annotation was not overlapping with a given DRegBC region. The total size of the eps1000 DRegBC region was then added to only negative value start and end flank numbers to give the total number of bp a given feature has within a given eps1000 DRegBC region. All remaining regions that did not have negative numbers added were then 100% contained within a given ChromHMM annotation. Any ChromHMM annotations that bordered with a clock region by only 1 bp were removed. The total bp annotation values within a eps1000 DRegBC region were then calculated as proportion within the total hyper- or hypomethylating region bp of the eps1000 DRegBC, i.e. each total feature bp in the hypermethylating regions were divided by the total number of bp of the hypermethylating clock regions and multiplied by 100 (the same procedure is done for hypomethylating annotations/regions). A list of 500 bp probes and eps1000 regions, both annotated with ChromHMM labels as well as calculated start/end flanks and bp overlap, can be found in **S.Table 5**.

To compare with approximate proportions of genomic chromatin states captured in RRBS, 500 bp probes were created for the Petkovich training data (CpGs *<*5 reads filtered out) using the “running window generator” function in SeqMonk. Read count was quantitated and regions with 0 reads were removed (hence only regions with CpGs *>*= 5 reads were retained). These probes were then annotated with ChromHMM labels in SeqMonk in the same manner as for the RegBCs. Overlap between 500 bp probes and ChromHMM were calculated in the same manner as the eps1000 DRegBC regions (see previous paragraph). Proportion of genomic chromatin states was then calculated as the total bp for each chromatin state divided by the total bp of chromatin overlapping with the 500 bp probes (204,291,600 bp), multiplied by 100. Next, for eps1000 DRegBC regions and 500 bp genomic regions, “Repressed” and “Heterochromatin” annotations were merged and labelled as “Heterochromatin”, with bp of both annotations added together, given their similar properties. Log2 fold change between the genomic chromatin proportions and the chromatin proportions in the DRegBC regions was calculated. The total bp of chromatin states for either hyper- or hypomethylating regions were divided by total bp of the corresponding chromatin state genomic bp total, then log2 of the resulting calculation was plotted using ggplot2. Any missing values prior to calculation (i.e. if the DRegBC regions did not have a particular ChromHMM annotation) were substituted with 0. Log2 of 0 values resulted in “Inf” which was substituted with 0 for the purposes of plotting. Calculated proportions and log fold change can be found in **S.Table 5**.

## Acknowledgements

We thank Chris Ponting, Riccardo Marioni and Neil Robertson for their valued feedback on the manuscript.

## Figures and legends

**Supplementary Table 1: Published bisuphite sequencing mouse clocks and the datasets used to train them**.

**Supplementary Table 2: Bisulphite sequencing mouse clocks applied to various datasets**. R^2^ and mean absolute error (MAE) values result from the clocks applied to all tissues in a given dataset.

**Supplementary Table 3: Various RegBCs and DRegBCs applied to training, cross-validation and external (Thompson et al. 2018) datasets**. Clocks highlighted bold are the regional clocks presented in Fig. 2.

**Supplementary Table 4: Annotation of 5kb RegBC and eps1000 DRegBC**.

**Supplementary Table 5: List of genomic 500 bp probes and eps1000 DRegBC regions annotated with ChromHMM states, as well as log fold change difference between genome and hyper/hypomethylating eps1000 DRegBC regions**.

## Bibliography

Andrews, Simon (2007). “Babraham Bioinformatics - SeqMonk Mapped Sequence Analysis Tool”. http://www.bioinformatics.babraham.ac.uk/projects/seqmonk/.

Andrews, Simon (2018). “PDF Manual: Analysing High Throughput Sequencing Data with SeqMonk”, p. 45.

Arneson, Adriana et al. (Dec. 2022). “A mammalian methylation array for profiling methylation levels at conserved sequences”. Nature communications 13.1.

Baheti, Saurabh et al. (Dec. 2016). “Targeted alignment and end repair elimination increase alignment and methylation measure accuracy for reduced representation bisulfite sequencing data”. BMC Genomics 17.1, p. 149.

Baker, George T. and Richard L. Sprott (Jan. 1988). “Biomarkers of aging”. Experimental Gerontology 23.4-5, pp. 223–239.

Bocklandt, Sven et al. (2011). “Epigenetic predictor of age”. PLoS ONE.

Bollati, Valentina et al. (Apr. 2009). “Decline in genomic DNA methylation through aging in a cohort of elderly subjects”. Mechanisms of Ageing and Development 130.4, pp. 234–239.

Bolzer, Andreas et al. (Ma. 2005). “Three-Dimensional Maps of All Chromosomes in Human Male Fibroblast Nuclei and Prometaphase Rosettes”. PLOS Biology 3.5, e157.

Cannon, Matthew V. et al. (Mar. 2014). “Maternal nutrition induces pervasive gene expression changes but no detectable DNA methylation differences in the liver of adult offspring”. PLoS ONE 9.3.

Chandra, Tamir et al. (2015). “Global reorganization of the nuclear landscape in senescent cells”. Cell Reports 10.4, pp. 471–483.

Chen, Brian H. et al. (2016). “DNA methylation-based measures of biological age: Meta-analysis predicting time to death”. Aging 8.9, pp. 1844–1865.

Christensen, Brock C. et al. (Aug. 2009). “Aging and Environmental Exposures Alter Tissue-Specific DNA Methylation Dependent upon CpG Island Context”. PLOS Genetics 5.8, e1000602.

Ernst, Jason and Manolis Kellis (Feb. 2012). “ChromHMM: automating chromatin-state discovery and characterization”. Nature Methods 2012 9:3 9.3, pp. 215– 216.

Ester, Martin et al. (1996). “A density-based algorithm for discovering clusters in large spatial databases with noise”. KDD 96.34, pp. 226–231.

Federico, C. Concetta et al. (Apr. 2006). “Gene-rich and gene-poor chromosomal regions have different locations in the interphase nuclei of cold-blooded vertebrates”. Chromosoma 115.2, pp. 123–128.

Federico, Concetta et al. (Dec. 2004). “The pig genome: compositional analysis and identification of the gene-richest regions in chromosomes and nuclei”. Gene 343.2, pp. 245–251.

Field, Adam E et al. (2018). “DNA Methylation Clocks in Aging: Categories, Causes, and Consequences”. Molecular Cell Review.

Fiorito, Giovanni et al. (Sept. 2021). “DNA methylation-based biomarkers of aging were slowed down in a two-year diet and physical activity intervention trial: the DAMA study”. Aging Cell 00, e13439.

Friedman, Jerome, Trevor Hastie, and Rob Tibshirani (2010). “Regularization paths for generalized linear models via coordinate descent”. Journal of Statistical Software.

Gilbert, Nick et al. (Sept. 2004). “Chromatin Architecture of the Human Genome: Gene-Rich Domains Are Enriched in Open Chromatin Fibers”. Cell 118.5, pp. 555–566.

Gopalan, Shyamalika et al. (Jul. 2017). “Trends in DNA methylation with age replicate across diverse human populations”. Genetics 206.3, pp. 1659–1674.

Gravina, Silvia et al. (Dec. 2016). “Single-cell genome-wide bisulfite sequencing uncovers extensive heterogeneity in the mouse liver methylome”. Genome Biology 17.1, p. 150.

Gu, Hongcang et al. (Mar. 2011). “Preparation of reduced representation bisulfite sequencing libraries for genome-scale DNA methylation profiling”. Nature Protocols 6.4, pp. 468–481.

Hahsler, Michael, Matthew Piekenbrock, and Derek Doran (2019). “Dbscan: Fast density-based clustering with R”. Journal of Statistical Software 91.

Han, Yang et al. (Aug. 2018). “Epigenetic age-predictor for mice based on three CpG sites”. eLife 7.

Han, Yang et al. (Dec. 2020). “Targeted methods for epigenetic age predictions in mice”. Scientific Reports 10.1, pp. 1–10.

Hannum, Gregory et al. (2013). “Genome-wide Methylation Profiles Reveal Quantitative Views of Human Aging Rates”. Molecular Cell.

Hepp, Tobias et al. (Ma. 2016). “Approaches to regularized regression - A comparison between gradient boosting and the lasso”. Methods of Information in Medicine 55.5, pp. 422–430.

Heyn, Holger et al. (Jun. 2012). “Distinct DNA methylomes of newborns and centenarians”. Proceedings of the National Academy of Sciences of the United States of America 109.26, pp. 10522–10527.

Higgins-Chen, Albert T. et al. (Aug. 2020). “Schizophrenia and Epigenetic Aging Biomarkers: Increased Mortality, Reduced Cancer Risk, and Unique Clozapine Effects”. Biological Psychiatry 88.3, pp. 224–235.

Horvath, Steve (2013). “DNA methylation age of human tissues and cell types”. Genome Biology 14.3156.

Horvath, Steve (2015). “Erratum to DNA methylation age of human tissues and cell types [Genome Biology, 14, R115, (2013)]”. Genome Biology 16.1, pp. 1–5.

Horvath, Steve and Andrew J. Levine (Nov. 2015). “HIV-1 Infection Accelerates Age According to the Epigenetic Clock”. Journal of Infectious Diseases 212.10, pp. 1563–1573.

Horvath, Steve et al. (Oct. 2014). “Obesity accelerates epigenetic aging of human liver”. Proceedings of the National Academy of Sciences of the United States of America 111.43, pp. 15538–15543.

Horvath, Steve et al. (Jul. 2016). “Huntington’s disease accelerates epigenetic aging of human brain and disrupts DNA methylation levels”. Aging 8.7, pp. 1485–1512.

Horvath, Steve et al. (Jul. 2018). “Epigenetic clock for skin and blood cells applied to Hutchinson Gilford Progeria Syndrome and ex vivo studies”. Aging 10.7, pp. 1758–1775.

Illingworth, Robert S. et al. (Sept. 2010). “Orphan CpG Islands Identify Numerous Conserved Promoters in the Mammalian Genome”. PLOS Genetics 6.9, e1001134.

Jansen, Rick et al. (Feb. 2021). “An integrative study of five biological clocks in somatic and mental health”. eLife 10, pp. 1–20.

Johansson, Åsa, Stefan Enroth, and Ulf Gyllensten (Jun. 2013). “Continuous Aging of the Human DNA Methylome Throughout the Human Lifespan”. PLoS ONE 8.6, p. 67378.

Kim, Youjin et al. (Jan. 2022). “Higher diet quality relates to decelerated epigenetic aging”. The American journal of clinical nutrition 115.1, pp. 163–170.

Klutstein, Michael et al. (Feb. 2017). “Contribution of epigenetic mechanisms to variation in cancer risk among tissues”. Proceedings of the National Academy of Sciences of the United States of America 114.9, pp. 2230–2234.

Koch, Carmen M. and Wolfgang Wagner (2011). “Epigenetic-aging-signature to determine age in different tissues”. Aging.

Krueger, Felix (2012). “Babraham Bioinformatics - Trim Galore!”

Krueger, Felix (2016). “Bismark Bisulfite Mapper-User Guide-v0.15.0”. http://bowtie-bio.sourceforge.net/bowtie2.

Krueger, Felix and Simon R. Andrews (Jun. 2011). “Bismark: A flexible aligner and methylation caller for Bisulfite-Seq applications”. Bioinformatics 27.11, pp. 1571–1572.

Kuhn, Max (Mar. 2020). “Classification and Regression Training [R package caret version 6.0-86]”.

Levine, Morgan E. et al. (Apr. 2018). “An epigenetic biomarker of aging for lifespan and healthspan”. Aging 10.4, pp. 573–591.

Lövkvist, Cecilia et al. (Jun. 2016). “DNA methylation in human epigenomes depends on local topology of CpG sites”. Nucleic Acids Research 44.11, pp. 5123–5132.

Lu, Ake T. et al. (Jan. 2019). “DNA methylation GrimAge strongly predicts lifespan and healthspan”. Aging 11.2, pp. 303–327.

Maegawa, Shinji et al. (Mar. 2010). “Widespread and tissue specific age-related DNA methylation changes in mice”. Genome Research 20.3, pp. 332–340.

Maierhofer, Anna et al. (2017). “Accelerated epigenetic aging in Werner syndrome”. Aging 9.4, pp. 1143–1152.

Marioni, Riccardo E. et al. (Dec. 2015). “DNA methylation age of blood predicts all-cause mortality in later life”. Genome Biology 16.1, p. 25.

Martin-Herranz, Daniel E. et al. (Aug. 2019). “Screening for genes that accelerate the epigenetic aging clock in humans reveals a role for the H3K36 methyltransferase NSD1”. Genome Biology 20.1, pp. 1–19.

McCormick, Helen et al. (Dec. 2017). “Isogenic mice exhibit sexually-dimorphic DNA methylation patterns across multiple tissues”. BMC Genomics 18.1.

Meer, Margarita V et al. (Nov. 2018). “A whole lifespan mouse multi-tissue DNA methylation clock”. eLife 7.

Meissner, Alexander et al. (2005). “Reduced representation bisulfite sequencing for comparative high-resolution DNA methylation analysis”. Nucleic Acids Research 33.18, pp. 5868–5877.

Meissner, Alexander et al. (Aug. 2008). “Genome-scale DNA methylation maps of pluripotent and differentiated cells”. Nature 454.7205, pp. 766–770.

Meuleman, Wouter et al. (Feb. 2013). “Constitutive nuclear lamina–genome interactions are highly conserved and associated with A/T-rich sequence”. Genome Research 23.2, pp. 270–280.

Miles, Jeremy (Sept. 2014). “R Squared, Adjusted R Squared”. In: Wiley StatsRef: Statistics Reference Online. Chichester, UK: John Wiley & Sons, Ltd.

Moqri, Mahdi et al. (Jun. 2022). “PRC2 clock: a universal epigenetic biomarker of aging and rejuvenation”. bioRxiv, p. 2022.06.03.494609.

Morgan M et al. (2022). “Rsamtools: Binary alignment (BAM), FASTA, variant call (BCF), and tabix file import. R package version 1.36.1”. https://bioconductor.org/packages/Rsamtools.

Olova, Nelly et al. (Mar. 2018). “Comparison of whole-genome bisulfite sequencing library preparation strategies identifies sources of biases affecting DNA methylation data”. Genome Biology 19.1, p. 33.

Orlanski, Shari et al. (Ma. 2016). “Tissue-specific DNA demethylation is required for proper B-cell differentiation and function”. Proceedings of the National Academy of Sciences of the United States of America 113.18, pp. 5018–5023.

Petkovich, Daniel A et al. (Apr. 2017). “Using DNA Methylation Profiling to Evaluate Biological Age and Longevity Interventions.” Cell metabolism 25.4, pp. 954–960.

Pintacuda, Greta et al. (Dec. 2017). “hnRNPK Recruits PCGF3/5-PRC1 to the Xist RNA B-Repeat to Establish Polycomb-Mediated Chromosomal Silencing”. Molecular Cell 68.5, pp. 955–969.

Rakyan, Vardhman K. et al. (Apr. 2010). “Human aging-associated DNA hypermethylation occurs preferentially at bivalent chromatin domains”. Genome Research 20.4, pp. 434–439.

Reizel, Yitzhak et al. (Ma. 2015). “Gender-specific postnatal demethylation and establishment of epigenetic memory.” Genes & development 29.9, pp. 923–33.

Saccone, Salvatore, Concetta Federico, and Giorgio Bernardi (Oct. 2002). “Localization of the gene-richest and the gene-poorest isochores in the interphase nuclei of mammals and birds”. Gene 300.1-2, pp. 169–178.

Simpkin, Andrew J. et al. (Jan. 2016). “Prenatal and early life influences on epigenetic age in children: A study of mother-offspring pairs from two cohort studies”. Human Molecular Genetics 25.1, pp. 191–201.

Simpson, Daniel J. and Tamir Chandra (Aug. 2021). “Epigenetic age prediction”. Aging Cell, e13452.

Simpson, Daniel J., Nelly N. Olova, and Tamir Chandra (Sept. 2021). “Cellular reprogramming and epigenetic rejuvenation”. Clinical Epigenetics 2021 13:1 13.1, pp. 1–10.

Slieker, Roderick C. et al. (Ma. 2018). “Age-related DNA methylation changes are tissue-specific with ELOVL2 promoter methylation as exception”. Epigenetics and Chromatin 11.1, p. 25.

Stubbs, Thomas M. et al. (Dec. 2017). “Multi-tissue DNA methylation age predictor in mouse”. Genome Biology 18.1, p. 68.

Thompson, Michael J et al. (Oct. 2018). “A multi-tissue full lifespan epigenetic clock for mice.” Aging 10.10, pp. 2832–2854.

Trapp, Alexandre, Csaba Kerepesi, and Vadim N. Gladyshev (Dec. 2021). “Profiling epigenetic age in single cells”. Nature Aging 2021 1:12 1.12, pp. 1189– 1201.

Wang, Tina et al. (Dec. 2017). “Epigenetic aging signatures in mice livers are slowed by dwarfism, calorie restriction and rapamycin treatment”. Genome Biology 18.1, p. 57.

Weidner, Carola I. et al. (Feb. 2014). “Aging of blood can be tracked by DNA methylation changes at just three CpG sites”. Genome Biology 15.2, R24.

Wickham, Hadley (2016). ggplot2: Elegant Graphics for Data Analysis.

Wu, Xiaohui et al. (Dec. 2019). “Effect of tobacco smoking on the epigenetic age of human respiratory organs”. Clinical Epigenetics 11.1, p. 183.

Zhang, Qian et al. (Dec. 2019). “Improved precision of epigenetic clock estimates across tissues and its implication for biological ageing”. Genome Medicine 11.1, p. 54.

Zhou, Wanding et al. (2022). “DNA Methylation Dynamics and Dysregulation Delineated by High-Throughput Profiling in the Mouse”.

